# Periodic variation of mutation rates in bacterial genomes associated with replication timing

**DOI:** 10.1101/195818

**Authors:** Marcus M. Dillon, Way Sung, Michael Lynch, Vaughn S. Cooper

## Abstract

The causes and consequences of spatiotemporal variation in mutation rates remains to be explored in nearly all organisms. Here we examine relationships between local mutation rates and replication timing in three bacterial species whose genomes have multiple chromosomes: *Vibrio fischeri, Vibrio cholerae*, and *Burkholderia cenocepacia*. Following five evolution experiments with these bacteria conducted in the near-absence of natural selection, the genomes of clones from each lineage were sequenced and analyzed to identify variation in mutation rates and spectra. In lineages lacking mismatch repair, base-substitution mutation rates vary in a mirrored wave-like pattern on opposing replichores of the large chromosome of *V. fischeri* and *V. cholerae*, where concurrently replicated regions experience similar base-substitution mutation rates. The base-substitution mutation rates on the small chromosome are less variable in both species but occur at similar rates as the concurrently replicated regions of the large chromosome. Neither nucleotide composition nor frequency of nucleotide motifs differed among regions experiencing high and low base-substitution rates, which along with the inferred ~800 Kb wave period suggests that the source of the periodicity is not sequence-specific but rather a systematic process related to the cell cycle. These results support the notion that base-substitution mutation rates are likely to vary systematically across many bacterial genomes, which exposes certain genes to elevated deleterious mutational load.

## IMPORTANCE

That mutation rates vary within bacteria genomes is well known, but the detailed study of these biases has been made possible only recently with contemporary sequencing methods. We applied these methods to understand how bacterial genomes with multiple chromosomes, like *Vibrio* and *Burkholderia*, might experience heterogeneous mutation rates because of their unusual replication and the greater genetic diversity found on smaller chromosomes. This study captured thousands of mutations and revealed wave like rate variation that is synchronized with replication timing and not explained by nucleotide content. The scale of this rate variation over hundreds of kilobases of DNA strongly suggests that a temporally regulated cellular process may generate wave-like variation in mutation risk. These findings add to our understanding of how mutation risk is distributed across bacterial and likely also eukaryotic genomes, owing to their highly conserved replication and repair machinery.

## INTRODUCTION

Mutation rates may vary within genomes for a variety of reasons, from straightforward causes like repetitive sequences causing polymerase slippage or the deamination and errant repair of methylated bases, to more complex causes like transcription-translation conflicts (1, 2). These processes tend to produce mutation rate heterogeneity over intervals less than 1 Kb. What is underappreciated is the potential for mutation rates to vary over longer ranges that may exceed 100 Kb and affect hundreds of genes. The prevalence and causes of long-range variation are unclear but have been attributed to effects of error prone polymerases (3), error prone repair pathways (4), and inconsistent nucleotide pools (5). If this long-range variation is common and systematic, the affected genes would be subject to greater mutational load and this process could select for gene reordering to avoid mutation risk.

On the other hand, replication timing, or the relative distance from the origin of replication, is one of the most conserved properties of orthologous genes, (6). Selection to maintain gene order has been attributed mostly to gene expression, where intragenic variation in the binding of nucleotide associated proteins (NAPs) and compaction of the nucleoid induce selection on gene order and location for optimal expression (6–9). Consequently, genes may face conflicts between the demand for optimal expression and their mutation risk, which has broad implications for genome evolution and genetic diseases. A series of comparative studies in multicellular eukaryotes (10–13), unicellular eukaryotes (12, 14), archaea (15), and bacteria (16, 17) have shown that synonymous substitution rates – a product of all population genetic forces including mutation, genetic drift, and selection – vary across the genome and generally increase in late replicating regions. This correlation could result from higher base-substitution mutation (bpsm) rates or weaker purifying selection in late replicating regions (1, 16, 18). A powerful approach to disentangle these processes is the mutation-accumulation (MA) experiment analyzed by whole-genome sequencing (WGS), in which many replicate lineages are passaged through hundreds of single cell bottlenecks in the near absence of natural selection and all mutations are identified. Our aim was to directly test whether *de novo* mutation rates vary among genome regions, and specifically whether such long-range systematic mutation rate variation operates in bacteria.

This study builds upon several prior MA-WGS studies in diverse bacterial species. Above all, mutation rates in bacteria are remarkably low, even dropping below 10^−3^/genome/generation (1, 19). Such low rates mean that MA experiments using wild-type strains with intact mismatch repair (MMR) fail to capture enough mutations to detect long-range mutation rate variation (19–21). MA studies with MMR-deficient organisms generate much larger collections of mutations but have shown no simple, linear correlation between bpsm rates and replication timing (19, 22–25). Thus, the more rapid evolution of late-replicated genes likely results from weaker purifying selection, not increased mutation rates. More intriguingly, MA studies of MMR-deficient bacteria, including *Escherichia coli*, *Pseudomonas fluorescens*, *Pseudomonas aeruginosa*, and *Bacillus subtilis* have revealed significant non-linear or periodic variation in mutation rates among genome regions (19, 22–25).

We chose to study three bacterial species with genomes containing multiple circular chromosomes: *Vibrio cholerae*, *Vibrio fischeri*, and *Burkholderia cenocepacia* (19, 21). This is an underappreciated but not uncommon bacterial genome architecture (16, 26–28) and enables effects of chromosome location and replication timing to be distinguished. Setting aside the distinction between chromosomes and megaplasmids (29), the *Vibrio cholerae* and *Vibrio fischeri* genomes are composed of two chromosomes, while the *Burkholderia cenocepacia* genome is composed of three. In each species, the first chromosome (chr1) is largest, harbors the most essential genes, and is expressed at the highest levels (16, 30). Secondary chromosomes (chr2, chr3) also initiate replication from a single origin and are replicated bi-directionally on two replichores (28, 31, 32). While they are replicated at the same rate as the first chromosome, their origins of replication (*oriCII*) have distinct initiation requirements from those of chr1 origins (*oriCI*) (26, 33). Importantly, chr2 (or chr3) replication is delayed relative to chr1 to ensure that replication of all chromosomes terminates synchronously (28, 32, 34). Consequently, the genome region near the origin of chr1 is always replicated prior to secondary chromosomes, while late replicated regions of chr1 are replicated concurrently with chr2.

This replication timing program in bacteria with multiple circular chromosomes enabled a test of whether secondary chromosomes experience similar mutation rates and regional variation to concurrently late replicated regions of primary chromosomes. Here we report detailed analyses of the genome-wide distribution of spontaneous bpsms generated by MA-WGS experiments with MMR-deficient strains of *V. fischeri* (4313 bpsms) and *V. cholerae* (1022 bpsms), and spontaneous bpsms generated by MA-WGS experiments with MMR-proficient strains of *V. fischeri* (219 bpsms), *V. cholerae* (138 bpsms), and *B. cenocepacia* (245 bpsms) (19, 21). We define the patterns of fluctuations in mutation rates within each genome and assess whether this variance affects coordinately replicated regions within and among chromosomes. In the MMR-deficient lines, we find evidence of systematic variation in mutation rate that implies that the causative factors act not just spatially but also temporally with the cell cycle, a phenomenon that could apply to a broad range of organisms.

## RESULTS

Two MMR-deficient (mutator) and three MMR-proficient (wild-type) MA-WGS experiments were founded by five different ancestral strains: a) *V. fischeri* ES114 *ΔmutS* (*Vf*-mut), b) *V. cholerae* 2740-80 *ΔmutS* (*Vc*-mut), c) *V. fischeri* ES114 wild-type (*Vf*-wt), d) *V. cholerae* 2740-80 wild-type (*Vc*-wt), and e) *B. cenocepacia* HI2424 wild-type (*Bc*-wt). Forty-eight independent MA lineages were propagated for 43 days in the two mutator experiments and seventy-five MA lineages were propagated for 217 days in the three wild-type experiments. In total, successful WGS was completed on evolved clones of 19 *Vf*-mut lineages, 22 *Vc*-mut lineages, 48 *Vf*-wt lineages, 49 *Vc*-wt lineages, and 47 *Bc*-wt lineages. Despite the fact that the mutator experiments were shorter and involved fewer lineages, the vast majority of bpsms were generated in the *Vf*-mut and *Vc*-mut lineages, as their bpsm rates are 317-fold and 85-fold greater than those of their wild-type counterparts, respectively. Consequently, effects of genomic position on bpsm rates can be studied in much greater detail in the mutator lineages, where adequate numbers of bpsms are distributed across the genome at intervals as short as 10 Kb (Table 1), the approximate length of bacterial microdomains (7).

**Table 1.** Number of base-substitution mutations (bpsms) in each mutation accumulation experiment, and the associated average number of bpsms in intervals of variable sizes.

| MA<br>Lines | No. of<br>bpsm | 500 Kb |  | 250 Kb |  | 100 Kb |  | 50 Kb |  | 25 Kb |  | 10 Kb |  |
| --- | --- | --- | --- | --- | --- | --- | --- | --- | --- | --- | --- | --- | --- |
|  |  | Avg. | SEM | Avg. | SEM | Avg. | SEM | Avg. | SEM | Avg. | SEM | Avg. | SEM |
| *Vf*-mut | 4313 | 499.00 | 35.82 | 253.50 | 13.77 | 101.05 | 3.12 | 50.53 | 1.26 | 25.26 | 0.51 | 10.08 | 0.18 |
| *Vc*-mut | 1022 | 141.50 | 21.14 | 65.33 | 6.23 | 25.47 | 1.53 | 12.51 | 0.59 | 6.22 | 0.25 | 2.50 | 0.09 |
| *Vf*-wt | 219 | 22.25 | 3.28 | 12.25 | 1.39 | 4.95 | 0.40 | 2.48 | 0.20 | 1.24 | 0.09 | 0.50 | 0.03 |
| *Vc*-wt | 138 | 18.00 | 3.09 | 8.75 | 0.95 | 3.42 | 0.30 | 1.72 | 0.14 | 0.83 | 0.07 | 0.34 | 0.03 |
| *Bc*-wt | 245 | 15.90 | 1.43 | 7.58 | 0.57 | 3.27 | 0.22 | 1.62 | 0.11 | 0.81 | 0.05 | 0.32 | 0.02 |

In comparing the overall bpsm rates between chromosomes in the mutator lineages, we observed that the bpsm rate on chr1 and chr2 of *V. fischeri* were not statistically distinguishable (χ^2^ = 0.11, df = 1, p = 0.741), while the bpsm rate on chr1 of *V. cholerae* was slightly higher than the rate of chr2 (χ^2^ = 4.54, df = 1, p = 0.0331) (19). However, even in *V. cholerae*, the variation in bpsm rates was minimal between chromosomes and our data suggested that considerably greater variation may exist within chromosomes (19). To determine the effects of genomic position on bpsm rates on a finer scale, we analyzed bpsm rates among intervals of varying size (10-500Kb) extending bi-directionally from the *oriCI* as the replication forks proceed during replication. Rates on chr2 were analyzed using the same intervals as chr1 but according to the inferred replication timing of *oriCI* (Figure S1A). This enables direct comparisons between concurrently replicated intervals on both chromosomes. To illustrate how this analysis works, we plotted the patterns of bpsm rates from a recent *E. coli* mutator MA experiment where mutation rates were demonstrated to vary in a wave-like pattern that is mirrored on the two replichores of its singular circular chromosome (20, 22) (Figure S1B). If replication timing is responsible for this pattern, a hypothetical secondary chromosome in *E. coli* would be expected to mirror concurrently replicated (late replicating) regions on the primary chromosome (Figure S1B).

### Base-substitution mutation rates are wave-like on chr1 in mutator lines

Mutation rates were not uniformly distributed across 10-500Kb intervals on chr1 in either the *Vf*-mut or the *Vc*-mut MA experiments (Supplementary Data), but we could not reject the null hypothesis of uniform rates on chr2, which has lower variance in bpsm rates. Variation in bpsm rates on chr1 in both *Vf*-mut and *Vc*-mut experiments follows a wave like pattern that is mirrored on both replichores bi-directionally from the origin of replication (Figure 1A, B). This mirrored pattern is evident at multiple interval sizes and is consistent with what has been reported on the single chromosome of *E. coli* (22), although the length of the wave periods observed here are shorter (Figure S1B). The waveform of bpsm rates is low near *oriCI*, increases to its peak approximately 600 Kb from the *oriCI* on both replichores, and declines into another valley before rising and falling again in the approach to the replication terminus. Two distinct waves can be seen on each replichore of chr1 (Figure 1A, B) but are less evident on chr2 (Figure S2). We focused our most detailed analyses of patterns of bpsm rate variation at the 100 Kb interval because it maximizes bpsms/interval while retaining two apparently mirrored bpsm rate waves on each replichore. Over 100Kb intervals, we see a significantly positive correlation between bpsm rates of concurrently replicated regions on the left and right replichores of chr1 in both *V. fischeri* and *V. cholerae* (Figure 2A, B). This relationship is also significant at most other interval lengths (Supplementary Data), but we find no such relationship when comparing 100 Kb intervals on the left and right replichores of chr2 as a consequence of its lower variance in bpsm rate.

**Figure 1.**
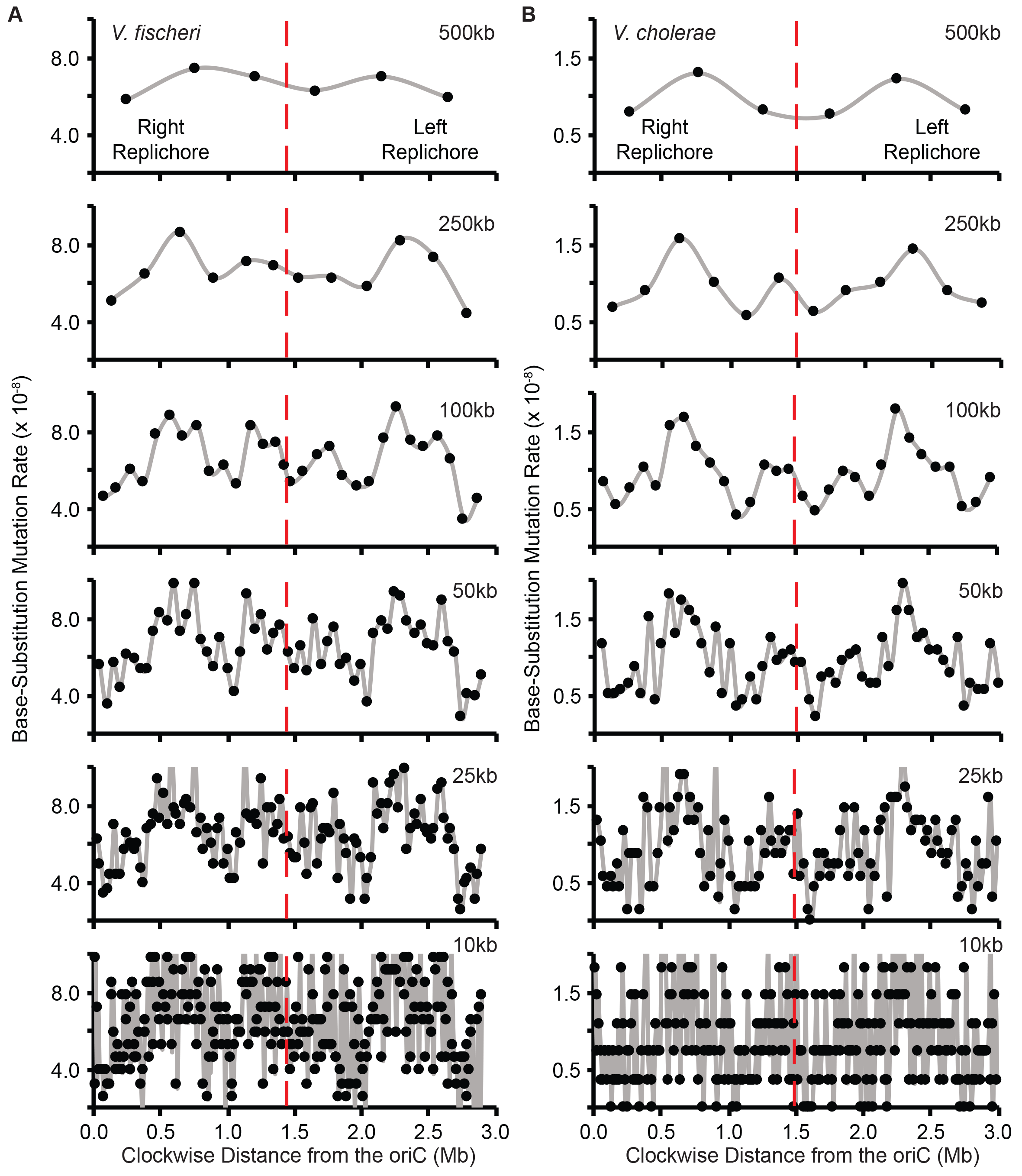
Patterns of base-substitution mutation (bpsm) rates at various size intervals extending clockwise from the origin of replication (*oriC*) in MMR-deficient mutation accumulation lineages of *V. fischeri* (A) and *V. cholerae* (B) on chromosome 1. Bpsm rates are calculated as the number of mutations observed within each interval, divided by the product of the total number of sites analyzed within that interval across all lines and the number of generations of mutation accumulation. The two intervals that meet at the terminus of replication (dotted red line) on each replichore are shorter than the interval length for that analysis, because the size of chromosome 1 is never exactly divisible by the interval length.

**Figure 2.**
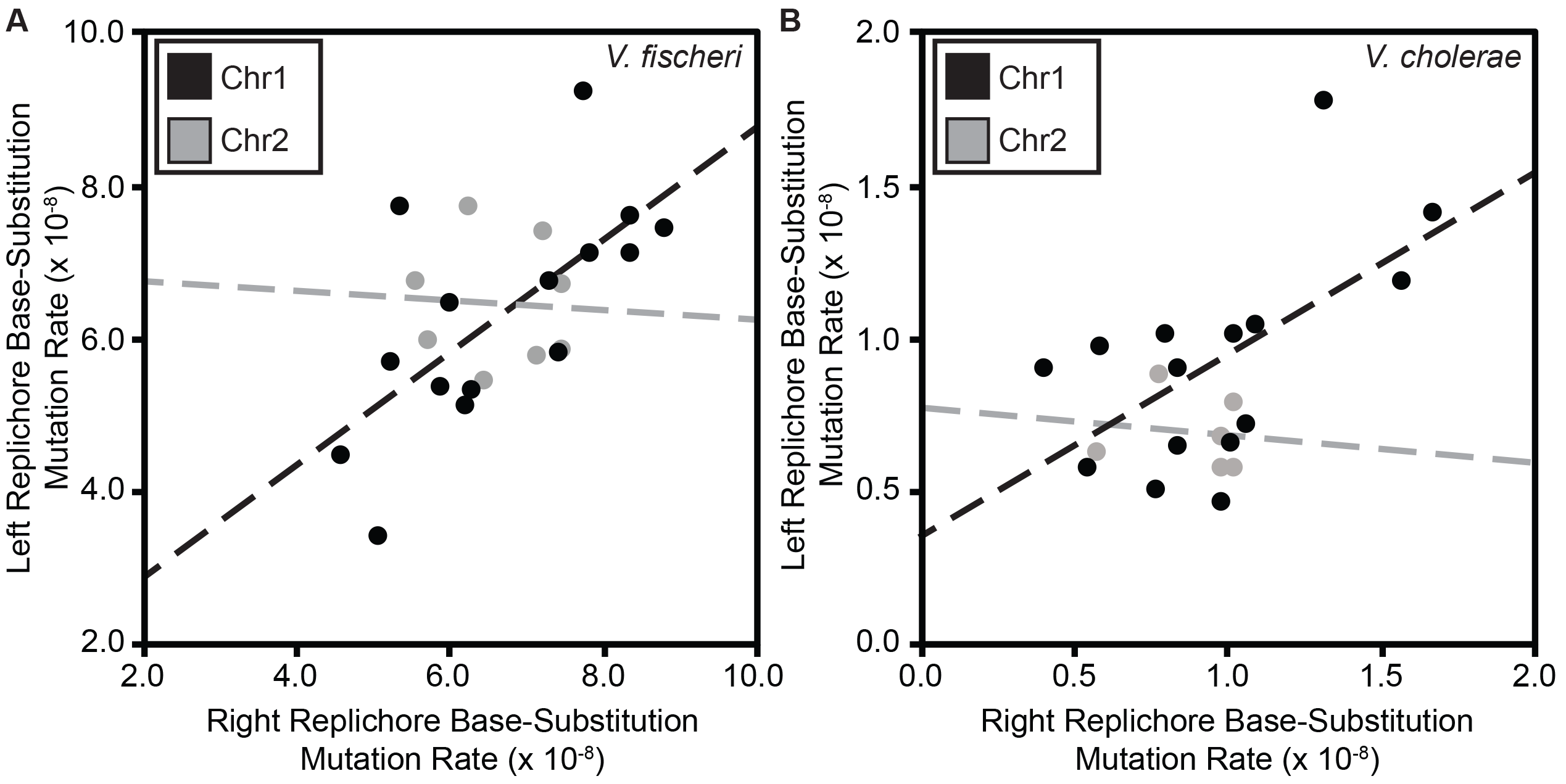
Relationship between base-substitution mutation (bpsm) rates in 100 Kb intervals on the right replichore with concurrently replicated 100 Kb intervals on the left replichore in MMR-deficient *Vibrio fischeri* (A) and *Vibrio cholerae* (B). Both linear regressions are significant on chr1 (V. *fischeri:* F = 10.98, df = 13, p = 0.0060, r^2^ = 0.46; *V. cholerae:* F = 6.76, df = 13, p = 0.0221, r^2^ = 0.34), but not on chr2 ( *V.fischeri:* F = 0.02, df = 6, p = 0.8910, r^2^ = 0.03 × 10^−1^; *V. cholerae:* F = 0.06, df = 4, p = 0.8140, r^2^ = 0.02).

### Concurrently replicated regions between chromosomes exhibit similar mutation rates

Given the observed relationship between bpsm rates of concurrently replicated regions on chr1, we might also expect late replicated regions of chr1 to experience similar bpsm rates as chr2 because of their concurrent replication. To study this relationship, we mapped the patterns of bpsm rates in 100 Kb intervals on chr2 to those of late replicated 100 Kb intervals on chr1 for both *Vf*-mut and *Vc*-mut (Figure S1). Fluctuations in bpsm rates on chr2 resemble those of late replicated regions on chr1 in both species (Figure 3A, B), but linear correlations in bpsm rates between chr1 and chr 2 were not significant (Supplementary Data). However, this lack of significant relationship may be a reflection of late replicated regions generally experiencing lower variance in bpsm rates than chr1 as a whole and given the strong resemblance in bpsm rate fluctuations between chr2 and concurrently late replicated regions of chr1, we attempted to falsify this match by correlating chr2 bpsm rates by correlating chr2 bpsm rates with all possible interval combinations on the right and left replichores of chr1. For *Vf*-mut, the lowest sum of the residuals (14.01 × 10^−8^) occurs when the chr2 intervals were mapped to the concurrently late replicated intervals on chr1 (Figure 3A; Supplementary Data). This same pattern was found for *Vc*-mut (Figure 3B; Supplementary Data). Thus, despite no significant linear correlation in mutation rate periodicity between chr1 and chr2, the spatial variation in bpsm rates on chr2 most closely resembles the rates of concurrently replicated regions on chr1 in both *V. cholerae* and *V. fischeri*. Interestingly, the delayed replication and small size of chr2 allows it to narrowly avoid the peak bpsm rates on the right and left replichores of chr1 in both *Vf*-mut and *Vc*-mut (Figure 3A, B). Thus, genes on chr2 may be subjected to less deleterious load than many of the genes on chr1, particularly in *V. cholerae*.

**Figure 3.**
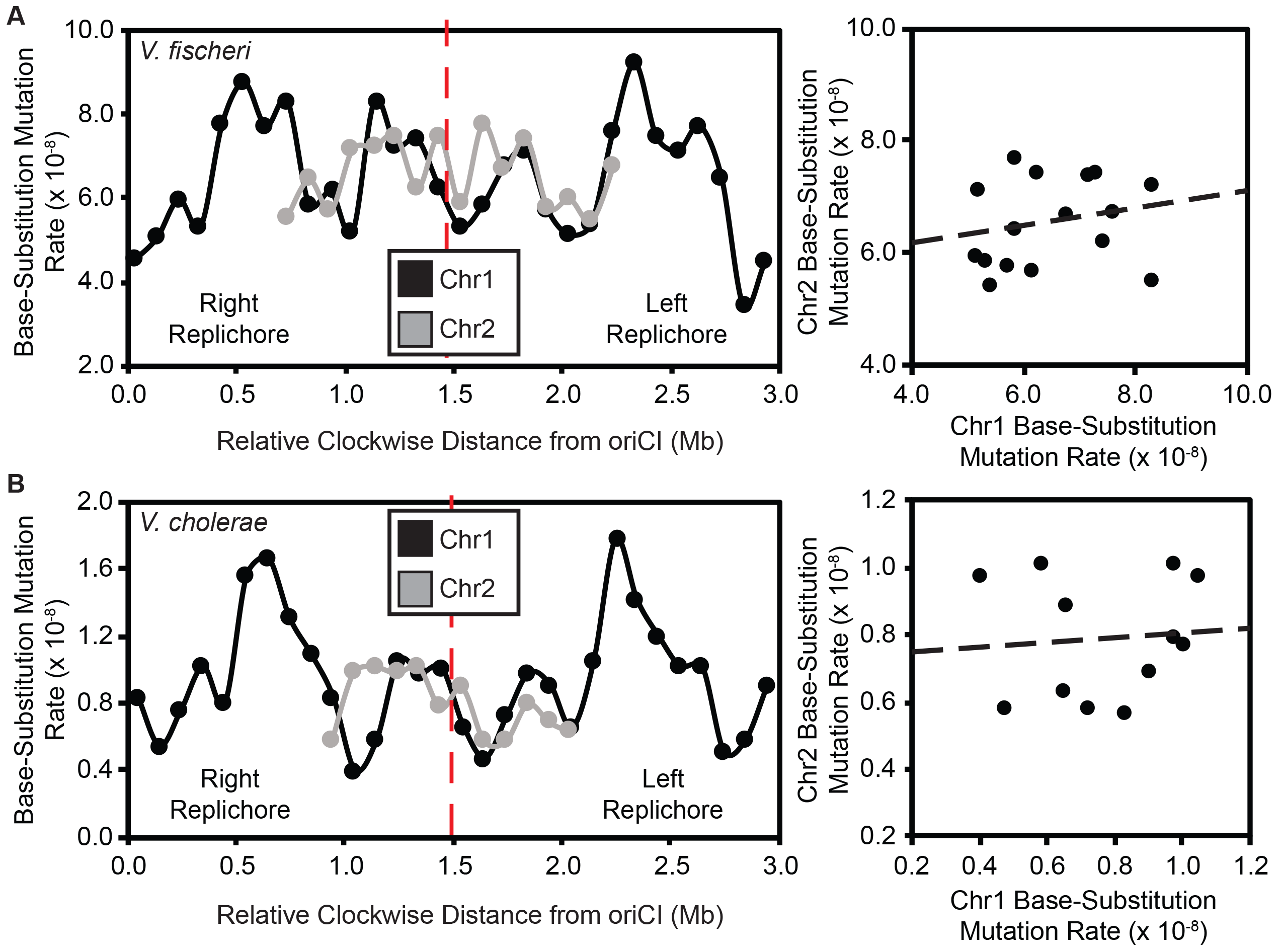
Patterns of base-substitution mutation (bpsm) rates in 100 Kb intervals extending clockwise from the origin of replication (*oriCI*) on chromosome 1 (chr1) and patterns of bpsm of concurrently replicated 100 Kb intervals on chromosome 2 (chr2) for MMR-deficient *Vibrio fischeri* (A) and *Vibrio cholerae* (B). Patterns of bpsm rates on chr2 appear to map to those of concurrently replicated regions on chr1 in both species, but the linear regressions between concurrently replicated intervals are not significant on chr1 and chr2 in either *V. fischeri* or *V. cholerae* (*V. fischeri*: F = 0.62, df = 14, p = 0.4442, r^2^ = 0.04; *V. cholerae:* F = 0.07, df = 10, p = 0.7941, r^2^ = 0.01).

### Wavelet transformations capture periodicities in base-substitution mutation rates

Recognizing that regional or cyclic variation in mutation rates may not be captured by linear models, we used wavelet transformations to characterize periodicities in the mirrored wave-like patterns in bpsm rates observed in this study. Bpsm rates on each chromosome in the *Vf*-mut and *Vc*-mut studies were transformed using the Morlet wavelet (35), which can reveal time-associated changes in the frequency of bpsms and has been successfully used in ecological time series analyses (36). This method was used to identify significant wave periods in bpsm rates on chr1 and chr2 and any variation in period length or amplitude across the chromosome. Significant wave periods of approximately 1.6 Mb and 0.8 Mb extend clockwise from *oriCI* in the *Vf*-mut lineages (Figure 4A). The single, long-period wave of 1.6 Mb is well supported across each replichore, while the shorter ~0.8 Mb period wave is significant across most of chr1 but its inferred length varies between 0.6 and 1.0 Mb. Thus, there are two synchronous periods per replichore or four periods in total around the chromosome, which are also clearly evident in Figure 1. These same two wave periods of approximately 1.6 Mb and 0.8 Mb were also ovserved in the bpsm rate data fond on chr1 in the *Vc*-mut lineages (Figure 5E).

**Figure 4.**
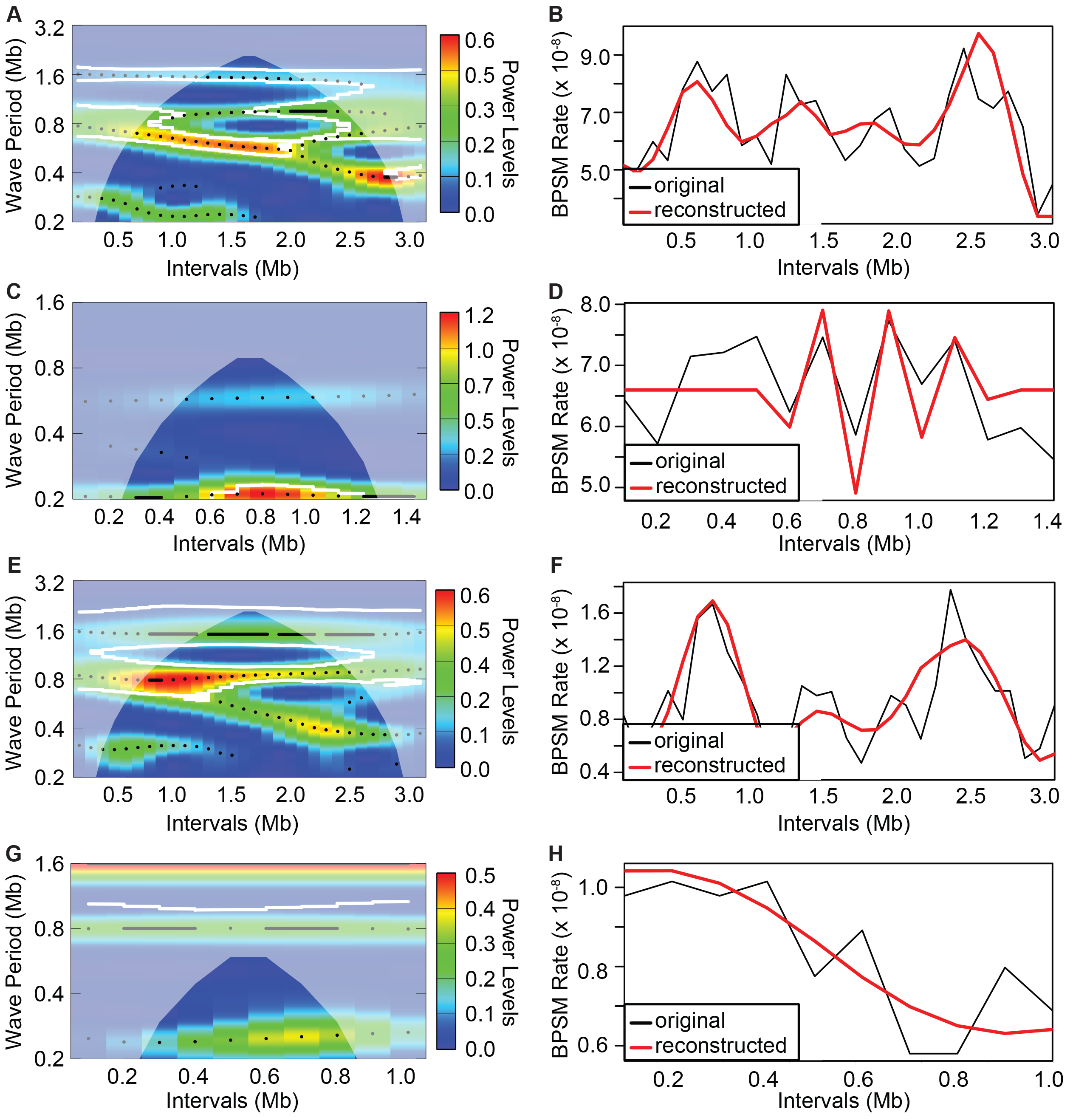
Wavelet power spectrum and resultant reconstruction of the patterns of base-substitution mutation (bpsm) rates in 100 Kb intervals extending clockwise from the *oriCI* region of chromosome 1 (A, B: *V. fischeri;* E, F: *V. cholerae*) and the *oriCII* region of chromosome 2 (C,D: *V. fischeri;* G,H: *V. cholerae)* using the MMR-deficient mutation accumulation lineages. White contour lines denote significance cutoff of 0.1 and wavelet power analyses follow an interval color key (A, C, E, G). Reconstructed series were generated using only the periods whose average power was significant over the entire interval (B, D, F, H).

**Figure 5.**
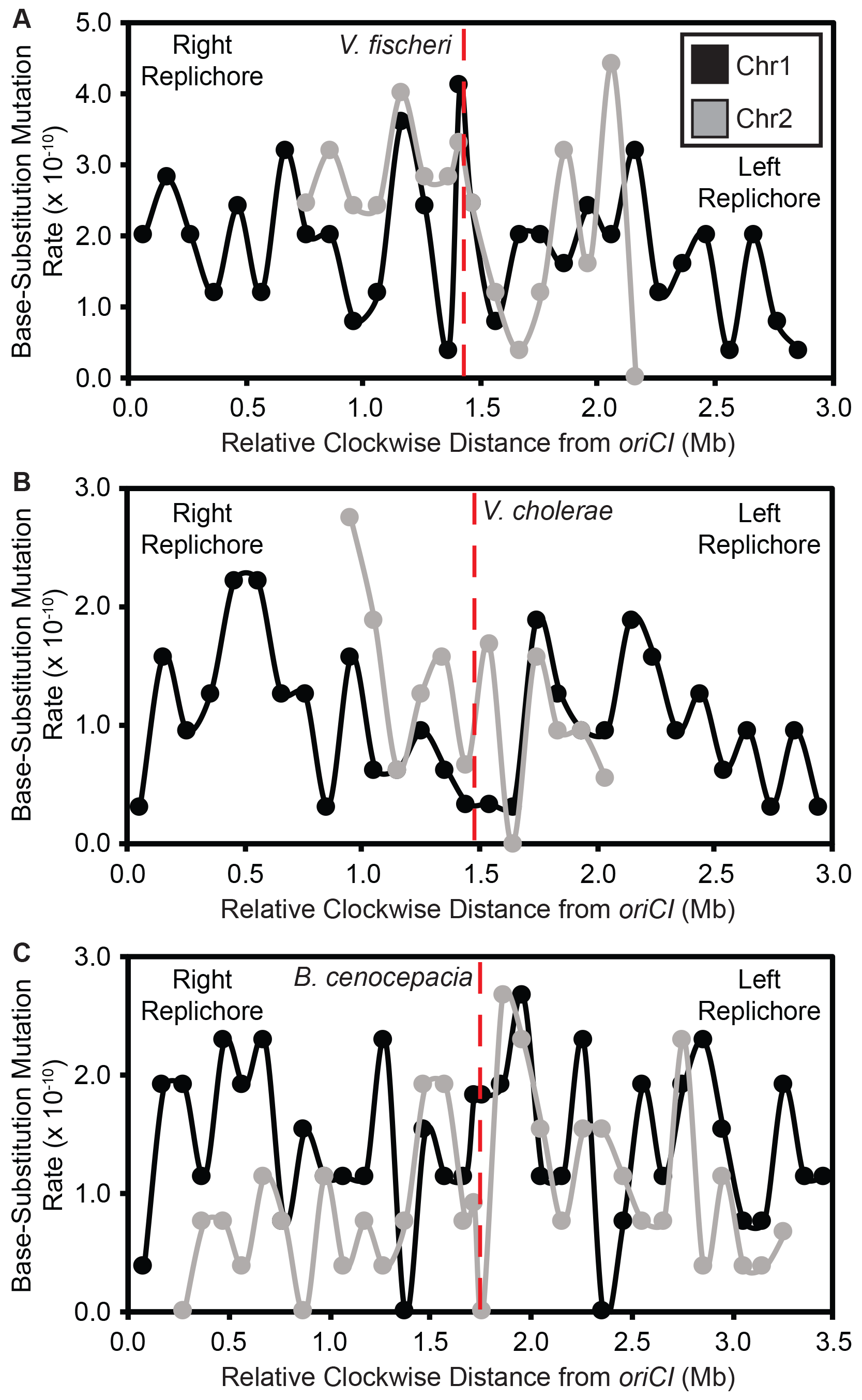
Patterns of base-substitution mutation (bpsm) rates in 100 Kb intervals extending clockwise from the origin of replication (*oriC*) on chromosome 1 (chr1) and concurrently replicated intervals of chromosome 2 (chr2) for WT (MMR+) *Vibrio fischeri* (A), *Vibrio cholerae* (B), and *Burkholderia cenocepacia* (C). *B. cenocepacia* also has a third chromosome, which is not shown. These visual patterns are not statistically significant, perhaps owing to low sample size: (linear regression; *Vf*-wt: F = 0.16, df = 14, p = 0.7001, r^2^ = 0.01; *Vc*-wt: F = 2.72, df = 10, p = 0.1300, r^2^ = 0.21; *Bc*-wt: F = 0.32, df = 30, p = 0.5760, r^2^ = 0.01)

Using only these wave models, we successfully reproduced the apparent periodicity of the 100 Kb data in both the *Vf*-mut and *Vc*-mut lineages (Figure 4B, F). Next, using the cross-wavelet transformation method to identify shared periodicities between replichores (35), we found that the wave model derived from one replichore predicts the behavior of the other (Figure S3A, B). It is also noteworthy that the statistically synchronous waves become smaller near the replication terminus, particularly in the *Vf-mut* lineages (Figure S3A, B), which is also apparent in the raw data presented in Figure 1. Perhaps because of this lower variation in late replicated regions, these modeling efforts were not successful on chr2 for either the *Vf*-mut or the *Vc*-mut experiment (Figure 4C, D, G, H).

### Replication-associated periodicity results from specific forms of base-substitution mutations

Nucleotide content varies across chromosomes and could conceivably underlie variation in bpsm rates among 100 Kb intervals. To address this possibility, we focused on A:T>G:C and G:C>A:T transitions in the *Vf*-mut and *Vc*-mut studies, as these two forms of bpsm represent 97.93% and 98.34% of all observed bpsms, respectively (19). Nucleotide composition did not vary significantly among 100 Kb intervals on chr1 or chr2. However, the spectra of bpsms corrected for nucleotide content varied significantly among intervals on chr1 in both the *Vf*-mut and *Vc*-mut MA experiments (Chi-square test; A:T>G:C: *Vf*-mut - χ^2^ = 62.26, df = 29, p = 0.0003, *Vc*-mut - χ^2^ = 49.04, df = 29, p = 0.0110; G:C>A:T: *Vf*-mut - χ^2^ = 120.69, df = 29, p < 0.0001, *Vc*-mut - χ^2^ = 111.19, df = 29, p < 0.0001). On chr2, only G:C>A:T substitutions in the *Vf*-mut study varied among intervals (χ^2^ = 26.81, df = 15, p = 0.0300). Interestingly, G:C>A:T mutation rates exhibit the greatest variation among chr1 intervals in both the *Vf*-mut and *Vc*-mut studies, and the positive correlations in bpsm rates on opposing replichores are driven largely by G:C>A:T, not A:T>G:C bpsms (Figure S4). The periodicity in bpsm rates in the *Vf-mut* and *Vc-mut* lines is therefore not caused by differences in nucleotide content but is predominantly caused by G:C>A:T transitions.

The immediate 5’ and 3’ nucleotide context of the mutated base can also influence rates and could conceivably lead to periodicity if trimers vary among intervals. Indeed, genome-wide bpsm rates in both the *Vf*-mut and *Vc*-mut studies vary more than 50-fold depending on the 5’ and 3’ bases flanking the site of the bpsm (Figure S5). This phenomenon has been found in several bacterial genomes and was found to be driven by sites neighboring G:C base pairs or dimers including alternating pyrimidine-purine and purine-pyrimidine nucleotides having significantly elevated mutation rates (24). However, the product of trimer abundance and specific mutation rates cannot explain the distribution of bpsms measured here on chr1 in either *V. cholerae* and *V. fischeri* (Figure S5).

### Low base-substitution mutation rates in wild-type lineages reveal modest regional variation

Despite conducting longer MA experiments (217 days vs 43 days) and sequencing more lineages (48 vs 22) derived from wild-type, MMR+ ancestors of *V. fischeri, V. cholerae*, and *B. cenocepacia*, considerably fewer bpsms accumulated in these lines than MMR-lines. Consequently, we cannot reject the null hypothesis that bpsms are uniformly distributed across chr1, chr2, and chr3 (for *Bc*) in the *Vf*-wt, *Vc*-wt, or *Bc*-wt MA experiments (Supplementary Data). Furthermore, coordinately replicated regions of chr1 and chr2 also did not exhibit correlated mutation rates, likely because of low sample sizes (Figure 5A, B, C). Only a mean of 4.65 (0.38), 3.29 (0.28), and 3.08 (0.22) (SEM) bpsms per 100 Kb interval were detected for the *Vf*-wt, *Vc*-wt, and *Bc*-wt MA lineages, respectively. Using effect size estimates derived from the significant patterns in MMR- lines (see Supplemental Methods), we estimate that the 132 mutations found on chr1 in the *Vf*-wt experiment would reveal a significantly non-uniform distribution of bpsms in only 19.46% of cases. The same analysis applied to the *Vc*-wt experiment predicts that significant regional variation in bpsms would be identified only 43.95% of the time. Further, applying effects from the *Vc*-mut to the *Bc*-wt experiment suggests that significant regional variation would be seen on chr1 in 55.16% of cases. Greater replication may be needed to capture more mutations in wild-type genomes to determine whether the periodicity in mutation rates seen in mutator lines also occurs in wild-type genomes, but we did observe that the patterns of bpsm rate variation in the *Vc*-wt experiment, where the effect size was largest, correlate with that of the corresponding mutator experiment, which implies a common underlying process for variation in mutator and wild-type bpsm rates (linear regression, 100 Kb intervals; *Vc*-wt - *Vc*-mut: F = 5.07, df = 38, p = 0.0303, r^2^ = 0.12).

## DISCUSSION

Variation in mutation rates among genome regions can have important implications for genome evolution and diseases, including most cancers (6–8, 37–40). One of the most conserved properties of genome organization is the relative distance of genes from the origin of replication (6, 41), which is expected to result in the long-term conservation of traits like expression and mutation rates for genes harbored in divergent genomes. Consequently, molecular modifications that change genome-wide patterns of replication timing, expression, and mutation rates could increase the probability of acquiring defective alleles in typically conserved regions, leading to disease. Indeed, alteration of the replication timing program can be an early step in carcinogenesis and a number of other somatic disease states (37). However, given the remarkable diversity in genome architecture across the tree of life, we still have much to learn about the nature of regional patterns of variation in bpsm rates and the genomic features and molecular processes that govern them.

Periodic variation in bpsm rates that is mirrored on the two replichores of bacterial chromosomes has been observed in genomes of some single-chromosome bacteria that are MMR-deficient (22, 25), yet not all species appear to experience this periodicity (23), and the underlying causes of periodic variation in bacterial bpsm rates are unknown. Here we demonstrate that MMR-deficient bacterial genomes with multiple chromosomes display mirrored, wave-like patterns of bpsm rates on chr1 (Figure 1A, B), and although we cannot reject the null hypothesis that bpsm rates are uniform on chr2, the patterns of bpsm rates on chr2 best match those of concurrently replicating regions on chr1 (Figure 3A, B). Furthermore, much of the genome-wide variation in bpsm rates that we observe appears to be generated by G:C>A:T transitions in both the *Vf*-mut and *Vc*-mut studies. Three MA experiments with MMR-proficient genomes hint at regional variation in bpsm rates, but these studies were insufficiently powered to reject the null hypothesis of uniformity. Nonetheless, shared periodicities in mutation rates between replichores and coarse similarities across chromosome regions that are coordinately replicated suggests strongly that mutation rates are affected by one or more common, global processes. Such a process influences replication fidelity throughout the genome at different active replication forks and causes bpsm rates to occur at a minimum level near the replication origin, rise to roughly 2-4 times these rates, then decline and repeat this cycle before replication termination. If physically separate genome regions share common mutation rates because of their shared replication timing, their genetic content may also be subject to common evolutionary forces.

This study cannot directly test the potential causes of mutation rate variation, but the bpsm patterns are more consistent with certain causes. First, nucleotide context can generate heterogeneous bpsm rates because certain nucleotides or nucleotide contexts are more prone to incur bpsms than others (20, 23, 24, 42, 43), and there is reason to believe that concurrently replicated regions on opposing replichores contain symmetrical gene content (41). Although we find that bpsm rates in both the *Vf*-mut and *Vc*-mut studies vary more than 50-fold depending on the bases flanking the site of the bpsm (Figure S5), this variation cannot explain the overall rate periodicity.

The replication machinery itself may also generate heterogeneous bpsm rates because of biased usage of error prone polymerases (3) or repair pathways (4) in certain genome regions. Both mechanisms have been invoked to explain why substitution rates scale positively with replication timing (4, 10–17), but the majority of these studies were performed in eukaryotes, and it is difficult to imagine how they might create the mirrored wave-like patterns of bpsm rates observed in bacterial chromosomes across 100 Kb intervals. Indeed, a series of MA studies in *E. coli* have shown that error-prone polymerases have minimal effects on mutation rates in the absence of DNA damage or stress (44).

Other genomic features that vary systematically with replication timing like binding of nucleoid-associated proteins (NAPs), transcription levels, and compaction of the bacterial nucleoid are also candidates for explaining our observed patterns of bpsm rates (6–9, 45). Sigma factors, DNA gyrase, and a number of NAPs have mirrored patterns of activity on the right and left replichores of the single chromosome in *E. coli* (6), possibly resulting from their concurrent replication. The resultant negative DNA superhelicity does correlate positively with the mirrored wave-like patterns of bpsm rates on opposing replichores of *E. coli* (22) and patterns of extant sequence variation are significantly impacted by NAPs that bind the DNA at different growth phases (9). However, effects of NAPs on sequence variation among published genomes are relatively weak and unlikely to produce the 2-4 fold changes in bpsm rates observed across the long interval lengths used in this study (9). While transcription levels may also impact bpsm rates through gene expression and replication-transcription conflicts (46), oscillations in expression patterns and gene density are not consistent with concurrently replicated regions experiencing similar expression levels (47), and expression has not been significantly correlated with the patterns of bpsm rates in *E. coli* and other species (1, 22).

The G:C>A:T and G:C>T:A bpsm that drive much of the observed periodicity are consistent with damage induced by reactive oxygen species (ROS) (Figure S4). It is conceivable that the plate growth conditions in these MA experiments generate ROS and thus more oxidized bases such as O6-methylguanine (O^6^-meG) and 8-oxo-guanine (8-oxo-G) (48, 49). The O^6^-meG modification commonly results in G:C>A:T mutations, while the 8-oxo-G modification commonly results in G:C>T:A mutations. These mutations are typically corrected by MMR and thus should be more common in MMR deficient MA lines. It remains unclear how either the origin or failed repair of ROS-induced mutations would be periodic with respect to replication timing. Conceivably, early-replicated nucleotides on Chr1 might be repaired more frequently by alternative pathways like translesion synthesis (48) and/or access to these repair complexes might be diluted with each new round of replication. This hypothesis could be tested by MA-WGS experiments under conditions that alter ROS exposure (44). For example, one recent experiment that focused on how the antibiotic norfloxacin influenced mutation rates in *E. coli* also tested effects of added peroxide because antibiotics may kill by ROS (50). Remarkably, this study also found periodic mutation rates that were mirrored on both replichores in the peroxide-treated lines, but no periodicity was seen in the norfloxacin-treated lines (potentially because of slower growth), indicating that mutation-rate periodicity may be induced by cyclical ROS-mediated effects.

With these alternative explanations in mind, we suggest that the most straightforward dynamic that could produce wave-like bpsm rates is variation in levels of deoxyribonucleotides (dNTPs). We describe a simple model of how dNTPs per replication fork may vary with *Vibrio* replication in Figure 6. Synthesis of dNTPs is controlled by levels of ribonucleotide reductase (RNR), whose production is coordinated with the rate of DNA synthesis but reaches its maximum following the onset of DNA replication to meet demand (51, 52). High levels of dNTPs are mutagenic in many organisms because of increased probability of misincorporation (52–54). In slow-growing bacteria whose division rates exceed the time required for chromosome replication, dNTP availability should increase after the start of replication and transiently increase the mutation rate but then decline to a baseline (Figure 6A, B). This predicts that slow growth should cause no mutation-rate periodicity, as the results from antibiotic-limited *E. coli* MA lines suggest (50). However, when bacterial generation times are faster than the time required for chromosome replication, which is commonplace for fast-growing species like *E. coli* or *Vibrio*, new rounds of replication are initiated and proceed before the first round concludes (55). Multi-chromosome genomes like those of *Vibrio* species require the additional firing of *oriC2*, which generates another burst of dNTP synthesis (Figure 6C, D). Consequently, fast-growing bacteria may experience multiple pulses of elevated RNR activity as origins fire (51), but the mutational effects of successive pulses of dNTP synthesis should be diluted across a growing number of active replication forks. We suggest this dynamic can simply generate the wave-like bpsm pattern observed in these experiments (Figure 1, Figure 6) as well as those previously reported in *E. coli* (22). Importantly, subsequent rounds of overlapping replication of either chromosome would only marginally affect the basic periodicity because dNTPs are diluted across multiple replication forks. Further, the model may explain two key features of the waves observed in our MA experiments - the greater amplitude of the first wave nearer to the origin and the lower overall variance in mutation rates in late-replicated regions, which results from dNTP bursts being diluted across more active replication forks (Figure 3). This model may also explain why not all bacterial genomes appear to experience periodic mutation rates (23) if they grow more slowly than the time for chromosome replication. We acknowledge that this model is speculative and requires considerable additional study, although the associations between replication dynamics and RNR activity and dNTP pools and mutation rates are both well supported (53, 56–58). A related possibility is that this periodicity arises from imbalances between rNTP and dNTP pools, which has been demonstrated to be mutagenic (53, 59, 60). At a minimum, this simple model relating ribonucleotide availability with mutation-rate periodicity is empirically testable by additional MA-WGS with defined mutants and altered growth conditions.

**Figure 6.**
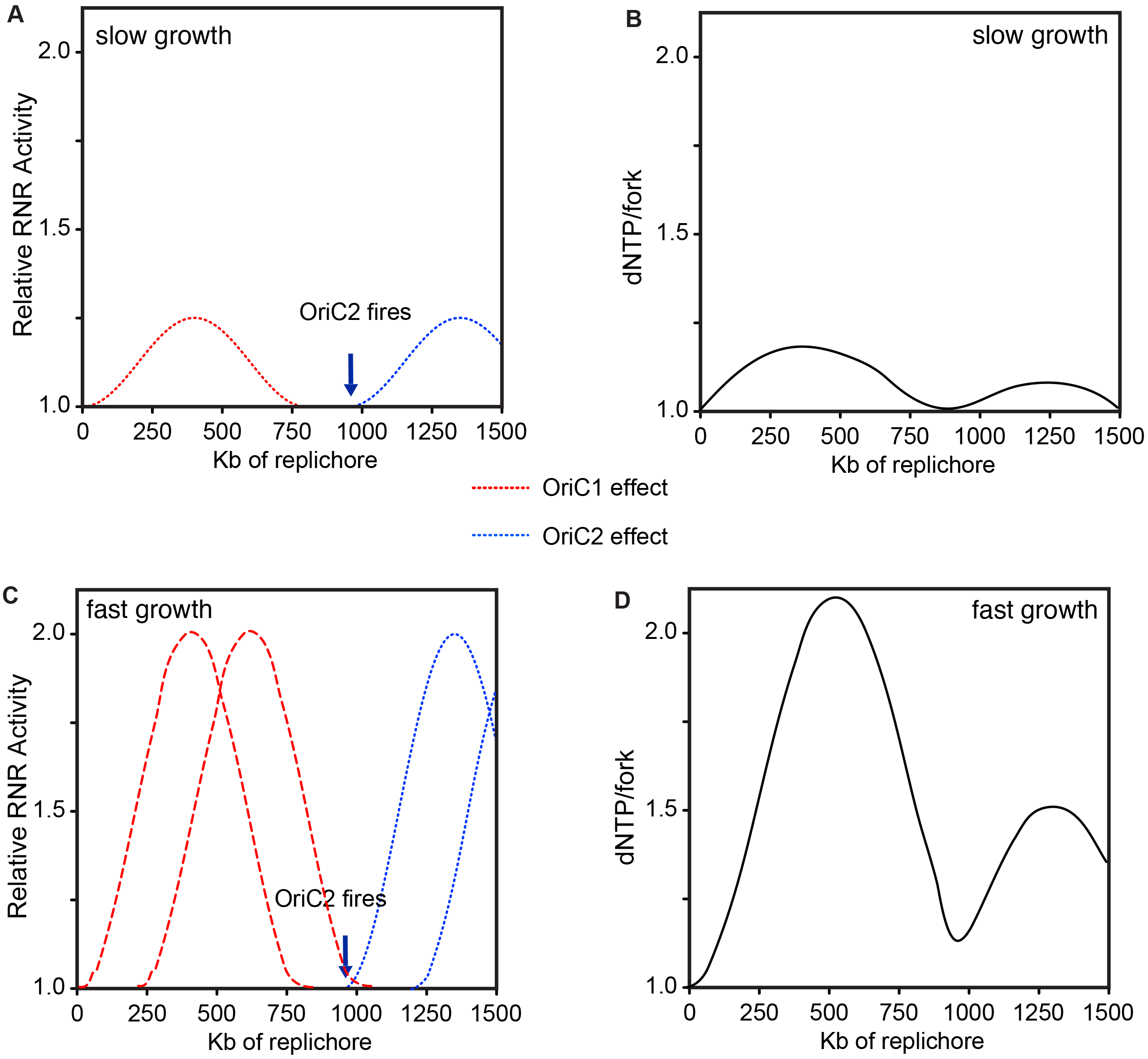
Hypothesized model of the relationship between replication timing, ribonucleotide reductase (RNR) activity, and the resulting availability of dNTPs per active replication fork. The model is fit to the *V. cholerae* genome with two chromosomes (chr) of 3.0 Mb and 1.1 Mb. RNR activity follows a wave that rises after the firing of the origin of chr1 and then steadily declines until additional origins fire. The chr2 origin should fire after ~950Kb of replication on each replichore of chr1 to ensure termination synchrony between chromosomes, stimulating a second wave of RNR activity. The right axis, in units of dNTP/fork, uses arbitrary relative units to depict how RNR activity is expected to increase dNTP pools to a maximum level (2.0) that is diluted by the number of concurrent, active forks. A,B: Under slow growth RNR activity rises and then falls to the baseline required to maintain synthesis. C,D: Faster growth requires a second round of replication. Note: further rounds of overlapping replication do not significantly alter predicted dNTPs/fork, the hypothesized driver of mutation-rate variability.

The conserved patterns of bpsm rates across concurrently replicated regions of MMR-lines also raises the question of whether these mutation biases influence the evolution of *Vibrio* genomes. In our previous studies of the mutation spectra from these experiments, higher rates of particular mutations were indeed found at synonymous sites among extant *Vibrio* and *Burkholderia* genomes (21, 61). If natural bpsm rates are in fact periodic in nature, we would expect genetic variation among strains to positively correlate with the bpsm rates in our defined 100 Kb intervals, particularly on chr1. We calculated the average pairwise synonymous (dS) and non-synonymous (dN) substitution rates in these intervals of *V. fischeri* and *V. cholerae* genomes (see Methods) and find a significant positive correlation for dS on chr1 but not on chr2 in *V. fischeri* (Figure S6). As expected from stronger selection on nonsynonymous sites, no significant correlation between dN and bpsm rates was found on either chromosome (Figure S6). No significant correlations between dS or dN and bpsm rates were found on either chromosome of *V. cholerae* (Figure S6). The scant correlations between evolutionary rates in coding sequences and spontaneous mutation rates may simply reflect that selection operating on both synonymous and non-synonymous sites is quite strong in bacteria (62). Alternatively, the natural patterns of bpsm rates in *V. fischeri* and *V. cholerae* may not be consistent with those observed in MMR- lines, which are strongly biased towards transition mutations. A more extensive study of the mutation spectra of wild-type genomes both experimentally and in natural isolates will determine the extent to which mutation-rate periodicity shapes genome evolution.

In summary, we have shown that bpsm rates in MMR-deficient lineages of *V. cholerae and V. fischeri* are non-uniformly distributed on chr1 and vary in a mirrored wave-like pattern that extends bi-directionally from the origin of replication. In contrast, late-replicated regions of chr1 and the entirety of chr2 experience more constant bpsm rates. These observations suggest that concurrently replicated regions of bacterial genomes experience similar bpsm rates prior to MMR, which could be governed by a number of temporally regulated cellular processes, including ROS, variation in dNTP pools, and the availability of replication machinery with secondary rounds of replication. We encourage research to disentangle effects of these cellular processes on bpsm rates (see for example (63)) as well as the signatures of these processes in natural populations, which will deepen our understanding of how mutation rates vary within genomes. Recalling that the relative distance of genes from the origin of replication is highly conserved across broad phylogenetic distances for a variety of functional reasons (6), it is quite possible that some genes are exposed to elevated mutational load while others are more shielded. In light of the growing effort towards evolutionary forecasting in microbial genomes (64), the need to determine whether the probability of new mutations substantively differs between genome regions is all the more pressing.

## METHODS

### Bacterial strains and culture conditions

MMR-deficient ancestors were generated by replacing the *mutS* gene in *V. fischeri* ES114 and *V. cholerae* 2740-80 with an erythromycin resistance cassette, as described previously (65–68). Complete genome sequences of these ancestors are publicly available (69, 70) or were generated by us for this project (71). Replication origins were determined using Ori-finder (19, 72, 73). MA experiments with *Vf*-mut and *Vf*-wt were conducted on tryptic soy agar (TSA) plates plus NaCl (30 g/liter tryptic soy broth powder, 20 g/liter NaCl, 15 g/liter agar) and incubated at 28°. MA experiments with *Vc*-mut, *Vc*-wt, and *Bc*-wt were conducted on TSA (30 g/liter tryptic soy broth powder, 15 g/liter agar) and incubated at 37°. MA experiments with MMR-lines involved 48 independent lineages founded from single colonies of *Vf-mutS* or *Vc-mutS* and were propagated daily for 43 days. MA experiments with WT lines involved 75 lineages founded from single colonies of *Vf, Vc, or Bc* and were propagated daily for 217 days (21, 61).

### Base-substitution mutation rate analysis at different genome intervals

Genomes were divided into intervals of 10 Kb, 25 Kb, 50 Kb, 100 Kb, 250 Kb, and 500 Kb, and bpsms were categorized by interval and location. On chr1, these intervals start at *oriCI* and extend bi-directionally to the replication terminus to mimic the progression of the two replication forks. Rates of bpsm were analyzed on secondary chromosomes similarly but intervals were measured relative to the initiation of replication of *oriCI* rather than to *oriCII* (Figure S1). This enables direct comparisons between concurrently replicated intervals on chr1 and chr2 based on established models of secondary chromosome replication timing in *V. cholerae* (28, 32, 34). Matched intervals of the same length were defined on each chromosome (n.b. chromosomes are not perfectly divisible by interval lengths so some intervals are shorter). Bpsm rates in each interval were calculated as the number of mutations observed in each interval, divided by the product of the total number of sites analyzed in that interval across all lines and the total number of generations of mutation accumulation, so rates in shorter intervals could be directly compared to the full-length intervals. For independent analyses of A:T>G:C and G:C>A:T mutations, bpsm rates were calculated as the number of mutations observed in each interval, divided by the product of the total number of sites in that interval that could lead to the bpsm being analyzed (A+T sites for A:T>G:C; G+C sites for G:C>A:T) and the total number of generations of mutation accumulation.

### Wavelet Transformations

We used the R package WaveletComp to evaluate properties of the wave-like patterns in bpsm rates in *Vf* and *Vc* and to test whether waves on opposing replichores were synchronous (35). The periodicity of bpsm rates on each chromosome of the *Vf*-mut and *Vc*-mut lineages at an interval length of 100 Kb was analyzed, treating each chromosome as a univariate series starting at the origin of replication and extending clockwise around the chromosome. WaveletComp uses the Morlet wavelet to transform the series of mutation rates then tests the null hypothesis of no periodicity for all combinations of intervals and periods (35). We performed this analysis using the “white.noise” method, with no smoothing, and a period range of 0.2 Mb to the entire length of the respective chromosomes. Default settings were used for all other parameters.

To test whether opposing replichores on Chr1 where synchronous, we used a cross-wavelet transformation (35) to test the null-hypothesis that no joint periodicity (synchronicity) exists among the two series as they traverse the primary chromosome in opposite directions. We used default settings but turned off smoothing and specified a period range of 0.2 Mb to the entire length of Chr1 in both *V. fischeri* and *V. cholerae*.

### Sequencing and Mutation Identification

Methods for genome sequencing, mutation identification, and evolutionary rate analyses are described in Supplementary text.

### Data Availability

Accession numbers for all of the whole-genome sequencing data produced by this study are PRJNA256340 for *V. fischeri*, PRJNA256339 for *V. cholerae*, and PRJNA326274 for *B. cenocepacia*.

## ACKNOWLEDGMENTS

We thank Cheryl Whistler, Randi Foxall, and Brian VanDam for technical support and Kevin Culligan, Greg Lang, Jeffrey Lawrence, and Pat Foster for helpful discussion. M.D., W.S., M.L., and V.C. designed the research; M.D., and W.S. performed the research; M.D. analyzed the data; and M.D., and V.C. wrote the paper. The authors declare no conflict of interest.

This work was supported by the Multidisciplinary University Research Initiative Award from the US Army Research Office (W911NF-09-1-0444 to ML, P. Foster, H. Tang, and S. Finkel); a National Science Foundation CAREER Award (DEB-0845851 to VSC), and NASA Astrobiology Institute CAN-7 NNA15BB04A to VSC.

## SUPPLEMENTAL MATERIAL

**Figure S1.** Design of the interval analysis used in this study to enable direct comparisons of base-substitution mutation (bpsm) rates of concurrently replicated regions on chromosome 1 (chr1) and chromosome 2 (chr2). A) For all multichromosome species analyzed in this study, secondary chromosomes are split at their origin of replication (*oriCII*), and mapped directly to concurrently replicated intervals in late replicating regions of chr1. All intervals on both chromosomes are thus relative to the initiation of replication of *oriCI*, and the boundaries of the intervals are consistent with their replication timing. B) Patterns of bpsm rates on the single chromosome of *Escherichia coli* MG1655 rph+ Δ*mutL*, derived from (20), show a wave-like mirrored pattern of bpsm rates on the two opposing replichores. If replication timing governs this pattern, a hypothetical secondary chromosome would be expected to mirror patterns of bpsm rates of late replicated regions on the primary chromosome.

**Figure S2**. Patterns of base-substitution mutation (bpsm) rates at various size intervals extending clockwise from the origin of replication (*oriCII*), in MMR-deficient mutation accumulation lineages of *Vibrio fischeri* (A) and *Vibrio cholerae* (B) on chromosome 2. All interval breakpoints are plotted relative to the initiation of replication of *oriCI* so that the boundaries of the intervals are at identical locations.

**Figure S3**. Cross-wavelet power spectrum plots comparing the patterns of base-substitution mutation (bpsm) rates in 100 Kb intervals extending clockwise from the *oriCI* region to those extending counterclockwise from the *oriCI* region in MMR-deficient mutation accumulation lineages of *Vibrio fischeri* (A) and *Vibrio cholerae* (B). Plots were generated using the WaveletComp package for Computational Wavelet Analysis in R, using an interval color key, 100 simulations, and significant synchronicity cutoffs of p < 0.1 for contour (white lines) and p < 0.05 for arrows. Colors represent the cross-wavelet power values at each interval in the genome for all possible wave periods, from dark blue (low power) to dark red (high power).

**Figure S4:** Relationship between base-substitution mutation (bpsm) rates in 100 Kb intervals on the right replichore with concurrently replicated 100 Kb intervals on the left replichore for A:T>G:C (A) and G:C>A:T (B) bpsms in MMR-deficient *Vibrio fischeri* and C) A:T>G:C and D) G:C>A:T bpsms in MMR-deficient *Vibrio cholerae*. Only the relationship between G:C>A:T bpsm rates of concurrently replicated regions on chr1 is significantly positive (*Vf*: A:T>G:C: Chr1 - F = 1.77, df = 13, p = 0.2067, r^2^ = 0.12, Chr2 - F = 3.26, df = 6, p = 0.1209, r^2^ = 0.35; G:C>A:T: Chr1 - F = 13.32, df = 13, p = 0.0029, r^2^ = 0.51, Chr2 - F = 0.17, df = 6, p = 0.6947, r^2^ = 0.03; *Vc*: A:T>G:C: Chr1 - F = 0.24, df = 13, p = 0.6313, r^2^ = 0.02, Chr2 - F = 1.74, df = 4, p = 0.2574, r^2^ = 0.30; G:C>A:T: Chr1 - F = 28.99, df = 13, p = 0.0001, r^2^ = 0.6904, Chr2 - F = 0.15, df = 4, p = 0.7209, r^2^ = 0.04).

**Figure S5.** Effects of nucleotide context (trimer content) on bpsm rates. A) Heatmap of the context dependent base-substitution mutation (bpsm) rates for the 64 possible trimer combinations based on their lagging strand orientation in MMR-deficient mutation accumulation lineages of *Vibrio fischeri* (A) and *Vibrio cholerae* (B). B) Patterns of base-substitution mutation (bpsm) rates in 100 Kb intervals extending clockwise from the origin of replication (*oriCI*) in MMR-deficient mutation accumulation lineages of *Vibrio fischeri* (A) and *Vibrio cholerae* (B). Observed patterns of bpsm rates (gray lines) on chromosome 1 (Chr1) and chromosome 2 (Chr2) are compared to the expected patterns of bpsm rates (blue lines) based on the trimer content of the interval. Bpsm rates differ significantly from expectations based on trimer content: Chi-square test; *Vf*-mut: Chr1 - χ^2^ = 137.24, df = 29, p < 0.0001, Chr2 - χ^2^ = 20.04, df = 15, p = 0.1703; *Vc*-mut: Chr1 - χ^2^ = 107.55, df = 29, p < 0.0001, Chr2 - χ^2^ = 14.87, df = 1, p = 0.1887)

**Figure S6**. Relationship between base-substitution mutation rates (bpsm) with average synonymous substitution rates (left panels) and average non-synonymous substitution rates (right panels) of genes. *V. fischeri*, top panel, *V. cholerae*, bottom panel, Average synonymous and non-synonymous substitution rates were calculated using the average rates of all one-to-one orthologs shared between *V. fischeri* ES114 and *V. fischeri* MJ11, or between *V. cholerae* 2740-80 and *V. cholerae* HE-16 within each 100 Kb interval. Synonymous and non-synonymous substitution rates for individual genes were calculated as described in (Yang and Nielsen 2000). In *V. fischeri*, only the relationship between bpsm rates and synonymous substitution rates on chromosome 1 is significant (A: Chr1 - F = 8.32, df = 28, p = 0.0080, r^2^ = 0.23, Chr2 - F = 0.56, df = 14, p = 0.4681, r^2^ = 0.04; B: Chr1 - F = 2.14, df = 28, p = 0.1554, r^2^ = 0.07, Chr2 - F = 0.03, df = 14, p = 0.8692, r^2^ = 0.02), and in *V. cholerae*, none are significant (C: Chr1 - F = 0.43, df = 28, p = 0.5186, r^2^ = 0.02, Chr2 - F = 0.49, df = 10, p = 0.5010, r^2^ = 0.05; D: Chr1 - F = 0.02, df = 28, p = 0.8897, r^2^ = 0.01 × 10^−1^, Chr2 - F = 0.01, df = 10, p = 0.9218, r^2^ = 0.01 × 10^−1^).

**Data Set S1.** Summary of all base-substitution mutations identified in each of the five mutation accumulation experiments carried out for this study, Chi-squared statistics of tests for uniform mutation rates, linear regression statistics for correlations between replichores and chromosomes, and residual fit of mutation rates to different chromosome intervals.

